# Genetic incompatibilities do not snowball in a demographic model of speciation

**DOI:** 10.1101/2021.02.23.432472

**Authors:** Carlos A. Maya-Lastra, Deren A. R. Eaton

## Abstract

Two populations evolving in isolation can accumulate genetic differences over time that cause incompatibilities in their hybrid offspring. These “Dobzhansky-Muller incompatibilities” (DMIs) are predicted to accumulate at a rate faster than linear as the number of incompatible gene interactions “snowballs”. Here we show that this snowball prediction is an artifact of two unrealistic modeling assumptions that stem from ignoring demography. We introduce a new alternative “demographic speciation model” in which the rate of DMI accumulation between populations is affected by the efficiency of purifying selection to remove incompatibilities that arise within populations. This model yields new testable predictions for understanding the tempo and mode of speciation based on population demographic parameters. A large-scale empirical analysis of bird and mammal datasets supports a unique prediction of our model: a negative relationship between effective population sizes and speciation rates. Our results challenge views of the snowball theory, and of ecological speciation models rooted in positive selection, showing instead that purifying selection may play a major role in mediating speciation rates.

## Introduction

Lineages vary in the rates at which they evolve barriers to reproduction, whether by behavioral, structural, or genetic mechanisms (*1–4*). Understanding factors underlying this variation is relevant for modeling rates of speciation, and ultimately for explaining and conserving global patterns of biodiversity (*5, 6*). Although many of the most rapid radiations are characterized by exceptional rates of evolution of reproductive isolating mechanisms – as with for example, African lake Cichlids (*7*) – these same lineages do not necessarily exhibit exceptional rates of evolution of DMIs (*8*). This disconnect, between the rate at which isolating mechanisms evolve and the strength of postzygotic incompatibilities, is an expectation of reinforcement theory (*9*), where substitutions reducing the fitness of hybrids, even if only by small effects, can cause selection for increased assortative mating long before complete hybrid sterility evolves. The difficulties involved with measuring and detecting small effect DMIs, however, hinders our ability to accurately estimate their rate of evolution.

The rate at which DMIs accumulate between species has been a subject of interest since Orr (*10*) first developed a mathematical framework for modeling their evolution. Orr’s snowball theory predicts that DMIs accumulate faster than linearly with time, as each new substitution increases the number of possible pairwise gene interactions that could become incompatible (Fig. 1A). Numerous empirical studies have examined the relationship between divergence times and various correlated measurements of DMI accumulation, such as hybrid viability, to test this prediction (*11–17*), but few have found support for the snowball model (*18*). Similarly, mixed support has been found among studies that have more directly counted the number of loci with large effects on hybrid sterility (*19–21*). The disagreement among these studies suggests that our existing theories may be insufficient to describe the diversity of rates observed in nature.

**Figure 1:**
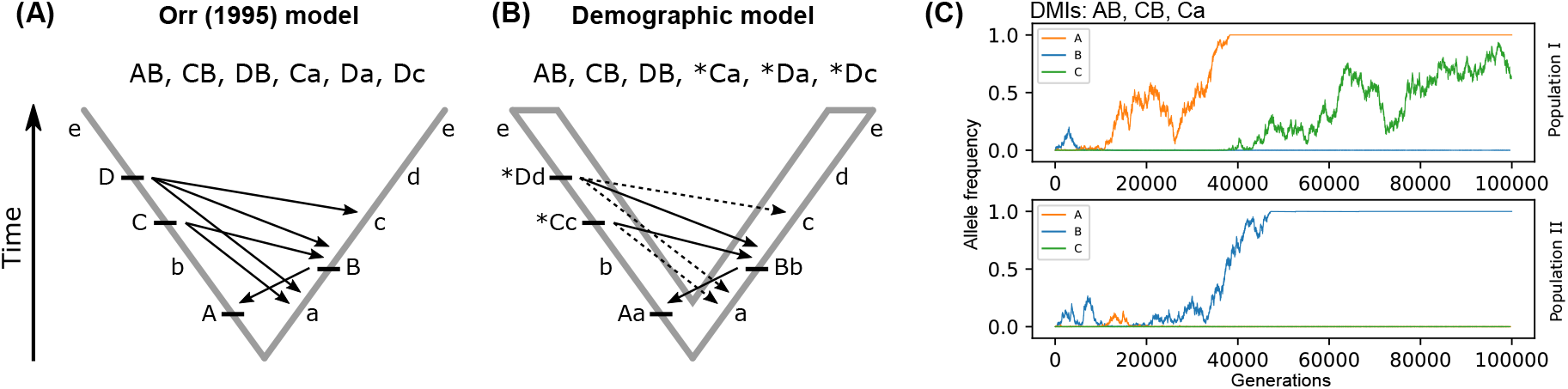
Theoretical predictions for the rate of accumulation of epistatic interactions causing Dobzhansky-Muller incompatibilities (DMIs) under (A) the Orr (1995) mathematical model and (B) our individual-based model. Arrows indicate incompatibilities between genes (labeled a-e) in two divergent populations. Derived mutations (uppercase) in either lineage may cause derived-derived incompatibilities (DDI; e.g., AB) or ancestral-derived incompatibilities (ADI; e.g., Ca). In Orr’s model derived mutations fix instantaneously, whereas in our model new mutations arise at low frequency and fluctuate by selection and drift. When new alleles do not fix instantly negative epistatic interactions can occur within parental populations (marked by *), affecting the fixation probability of alleles that would cause DMIs between species (shown as dashed arrows). (C) Purifying selection acts on the fitness consequences of negative epistatic interactions that arise among standing variation within populations, affecting the fixation probabilities of new alleles in each population (see description in text).

### The demographic speciation model

Here we revisit Orr’s snowball model and show that its faster-than-linear prediction is an artifact of two unrealistic modeling assumptions: the infinite-sites mutation model that does not allow for homoplasy, and the instant fixation of new mutations across all individuals in a population (i.e., no demography). While these simplifying assumptions have traditionally made it easier to model epistatic interactions, they obscure the effects of demography, selection, and the limits imposed by finite genome and population sizes.

We introduce a new individual-based model of DMI accumulation (demographic speciation model) that differs from Orr in allowing new mutations to evolve by drift and selection within two independently evolving finite populations with finite genomes (Fig. 1B). The absence of demography in Orr’s original model could be justified under the assumption that most DMIs are caused by substitutions that fix rapidly under positive selection. Our new model focuses instead on the much larger proportion of variation in genomes represented by neutral and nearly neutral deleterious mutations. This view aligns with our increasing knowledge of gene epistatic interaction networks (*22*), which considers that complex traits, such as hybrid fertility, are likely influenced by many small-effects originating from most or all genes in the genome.

An important consequence of modeling the evolution of epistatic interactions in a demographic framework is that standing genetic variation (the time to fixation of alleles) can expose incompatibilities not only in hybrids between populations, but also *within* the parental populations themselves (Fig. 1B). This introduces an opportunity for purifying selection to act on new combinations of alleles before they become fixed, which in turn can affect the rate of DMI accumulation between populations. The joint effects of demography and purifying selection on the accumulation of DMIs between species forms the basis of the demographic speciation model (Appendix I).

Purifying selection can remove genetic variants from a population based on their deleterious fitness effects, which in the context of our model, are defined by an allele’s epistatic interactions with other alleles in the genome. In this way, historical contingency plays a major role in mediating selection by affecting which alleles are present together in a population at any given time. This is demonstrated in an individual-based simulation for three genes under a demographic model with purifying selection where we define three potential DMIs that cause genetic incompatibilities (a deleterious fitness effect) in either population (Fig. 1C). In population I the derived allele “C” only increases to high frequency after allele “A” becomes fixed, because “Ca” incompatibilities cause selection against the “C” allele while the ancestral “a” is still present. In population II allele “B” instead increases to high frequency first, and consequently “A” and “C” both experience negative purifying selection. In each case, the order in which new alleles arise, and their frequencies, affects the fitness of subsequent alleles.

This simple realization has important consequences for modeling the evolution of DMIs between two independently evolving populations – notably, that they are not independent. Assuming that each lineage inherits the same ancestral network of gene interactions, they will also inherit the same set of ways in which the network can become disrupted (i.e., the same potential DMIs). This biases each lineage towards evolving along fewer possible ordered paths of fixation of new alleles (fixation-order paths). For example, if both populations in Fig. 1C had fixed the “A” substitution first, then each is more likely to evolve “C” next than “B”, since “AB” is incompatible. Similarly, neither is likely to have fixed “C” first, since it experiences negative epistasis with more alleles present in their common ancestor (e.g., “a” and “b”). Thus, in addition to affecting the probabilities of different fixation-order paths in each lineage, contingency (shared ancestry) also affects the probability that two lineages fix substitutions affecting the same gene interactions.

This expectation of the demographic speciation model stands in stark contrast to the snowball model. Whereas the infinite-sites mutational process in the snowball model makes homoplasy impossible, our model instead predicts that homoplasy – in the context of which sets of ancestral gene interactions are likely to be disrupted versus conserved – is a common feature among close relatives. If two populations fix derived substitutions in ways that preserve the same sets of ancestral gene interactions (i.e., perfect homoplasy) their F1 hybrids will exhibit zero DMIs; whereas if each population fixes substitutions affecting a non-overlapping set of gene interactions (i.e., infinite-sites model) they will exhibit the maximum possible number of DMIs in their hybrids. In this way, the extent of homoplasy in epistatic networks between two diverged populations can explain variation in their rate of DMI evolution.

The efficiency of purifying selection to remove deleterious variants will influence the extent to which two populations evolve along similar fixation-order paths. This is determined by the relative strength of selection versus genetic drift, and is mediated by population demographic parameters like effective population size (Ne). When Ne is high, each population is more likely to evolve along one of few potential fixation-order paths that minimize the evolution of deleterious epistasis within the population. When Ne is small, populations are more likely to evolve along one of many possible fixation-order paths that are more stochastic (dominated by genetic drift) and include the fixation of many slightly deleterious substitutions. Following from this logic, the extent of homoplasy between two diverged populations, and by extension, their rate of DMI evolution, should be predictable from population demographic parameters.

Here we use individual-based simulations to investigate the relationship between population demographic parameters, homoplasy in gene epistatic networks, and the evolution of DMIs between diverged populations. Our results provide theoretical and empirical evidence for the demographic speciation model, showing that by incorporating demography and purifying selection into models of speciation we can predict rates of DMI evolution that align better with empirical observations.

## Results

### The missing snowball

We used forward-in-time simulations (see Material and Methods) to explore seven models that vary in one or more assumptions that differentiate Orr’s snowball model (hereafter “Orr”) from an individual-based model (“Ind+Sel”; Table S1) representing our theoretical demographic speciation model. Of these seven models, only the Orr model, which includes the two unrealistic assumptions on which the snowball theory is based (infinite-sites mutations and the instant-fixation of new alleles) supports a quadratic rate of accumulation of DMIs (Fig. 2A). All other models support a decreasing rate through time that is best approximated by a logistic (or negative quadratic) fit (Table 1).

**Figure 2:**
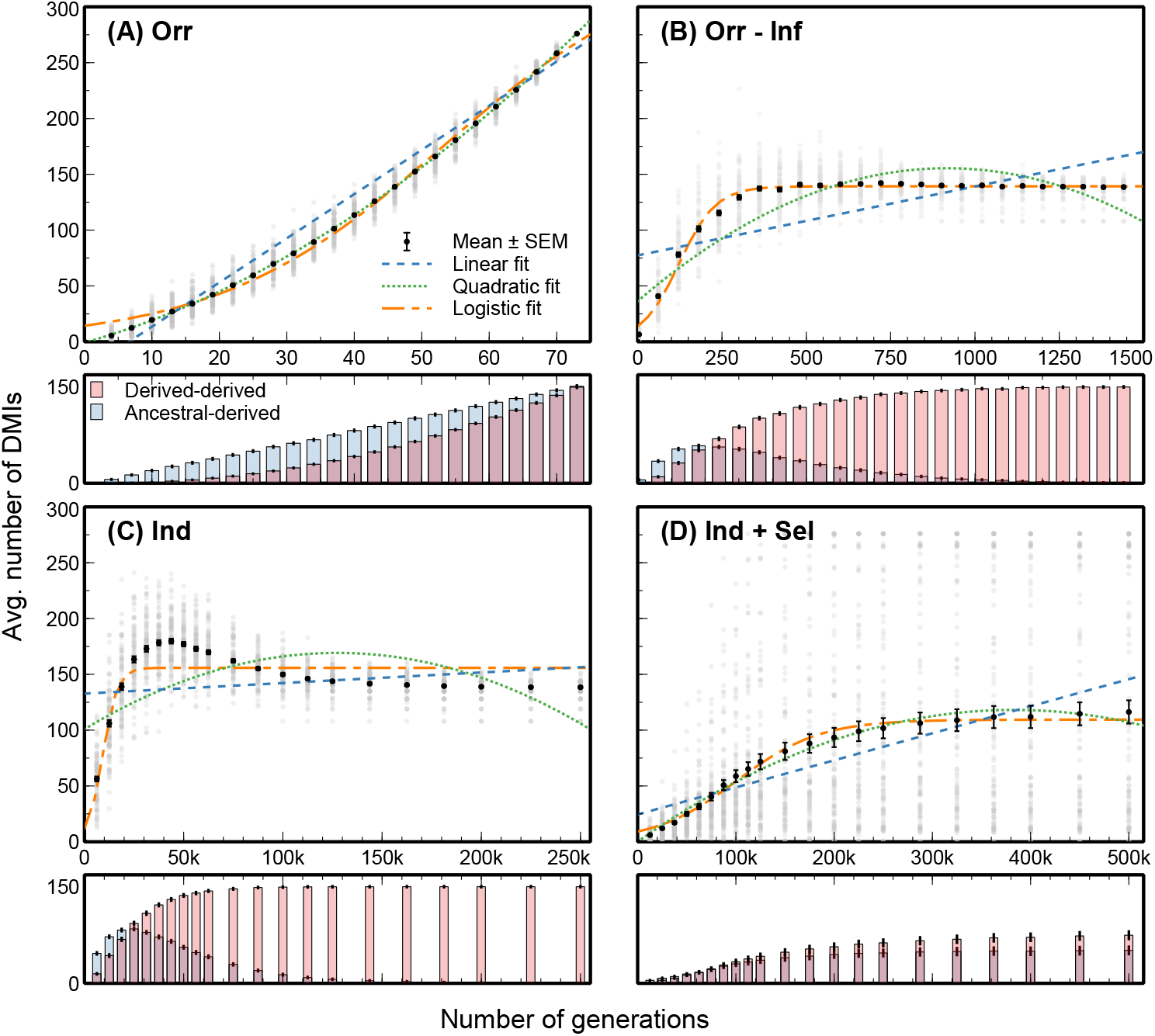
The rate of accumulation of DMIs in F1 hybrids between two diverged lineages in simulations from four models with different evolutionary assumptions. Only the Orr model (A) recovers the quadratic accumulation predicted by the snowball theory. All other models are best fit by a logistic function showing a slowdown in the rate of accumulation of DMIs through time (B-D). Lower panels decompose DMIs into DDI and ADI types, showing that ADI initially accumulate faster than DDI.

**Table 1.** Model comparison for fitting curves to the rate of DMI accumulation between two diverged species under seven evolutionary models that vary in their underlying assumptions. Only the ”Orr” model shows a quadratic, or even faster than linear, rate of accumulation of DMIs. All other models are best fit by a function in which the rate slows over time, either as a logistic or negative quadratic function (marked by *).

| Model | Assumptions |  |  |  | AICwt |  |  |
| --- | --- | --- | --- | --- | --- | --- | --- |
|  | Inst. | Inf. | Sel. | Rev. | Linear | Quadratic | Logistic |
| Orr | 1 | 1 | 0 | 0 | 0 | 1 | 0 |
| Orr-Inf | 1 | 0 | 0 | 0 | 0 | 0 | 1 |
| Orr-Inf+Rev | 1 | 0 | 0 | 1 | 0 | 0 | 1 |
| Ind | 0 | 0 | 0 | 0 | 0 | 0 | 1 |
| Ind+Rev | 0 | 0 | 0 | 1 | 0 | 0 | 1 |
| Ind+Sel | 0 | 0 | 1 | 0 | 0 | *0.14 | 0.86 |
| Ind+Rev+Sel | 0 | 0 | 1 | 1 | 0 | *1 | 0 |

By relaxing only the infinite-sites assumption of Orr’s model (Orr-Inf; Fig. 2B) the faster-than-linear pattern disappears, suggesting that the infinite-sites assumption underlies the snowball prediction. The fact that our results reproduce the expected snowball pattern when the Orr model assumptions are implemented confirms that other aspects of our simulation procedure – such as defining all possible DMIs before the simulation begins – do not artificially impact our results. While several studies have previously argued for incomplete support of the snowball theory and proposed different slowdown models (*18,26*), we argue that the snowball pattern is unlikely to evolve, since instead of predicting a probable range of rates under a realistic evolutionary scenario, it describes the maximum possible number of two-locus epistatic interactions under a single improbable mutational history.

To explain the slowing pattern of DMI accumulation observed in all other models we further decomposed epistatic interactions causing DMIs into two component types: derived-derived incompatibilities (DDI) and ancestral-derived incompatibilities (ADI). All models initially accumulate ADI faster than DDI, since a newly derived allele may be incompatible with many of the initially more common ancestral alleles. The Orr model is unique in that it always evolves to the maximum possible number of DMIs, while all other models evolve some degree of homoplasy that reduces DMIs. In the context of the simple binary-state gene model used here, homoplasy should be interpreted on the gene-functional level, in terms of the retention or loss of the ancestral gene function or interactions. An extreme example of homoplasy is apparent in neutral models (no selection) that relax the infinite-sites assumption (Orr-Inf and Ind), where the genome eventually becomes entirely saturated by derived states and all ADI are lost (Fig. 2B-C). This result, like the Orr model with its infinite-sites assumption, is unrealistic. In reality, many ancestral gene interactions should be retained, and this is the case when either reversibility (Fig. S1) or purifying selection (Fig. 2D) is included in the model. For this reason, we consider the individual-based model with purifying selection (Ind+Sel) to be the most realistic scenario for the evolution of DMIs in a demographic context.

### Purifying selection and parallel fixation orders

The evolution of homoplasy between the networks of gene interactions in two diverged lineages can be predicted by population demographic parameters. This is demonstrated in simulations from the Ind+Sel model, which applies purifying selection to individuals in each population based on the number of incompatible gene interactions in their genomes. This model yields two novel results: (*1*) high variance among replicate simulations in the rate and number of DMIs that evolve between populations (Fig. 2D); and (*2*) a negative correlation between the number of DMIs and effective population size (Fig. 3A). The first result is concordant with the high variance in rates of evolution of intrinsic post-zygotic reproductive isolation that has been reported in many empirical studies (*18*), and which the snowball model fails to predict. The second result, showing an effect of Ne on rates of DMI evolution, is only observed in models that include selection (Fig. S2), showing that it is a consequence of Ne’s effect on the strength of selection.

**Figure 3:**
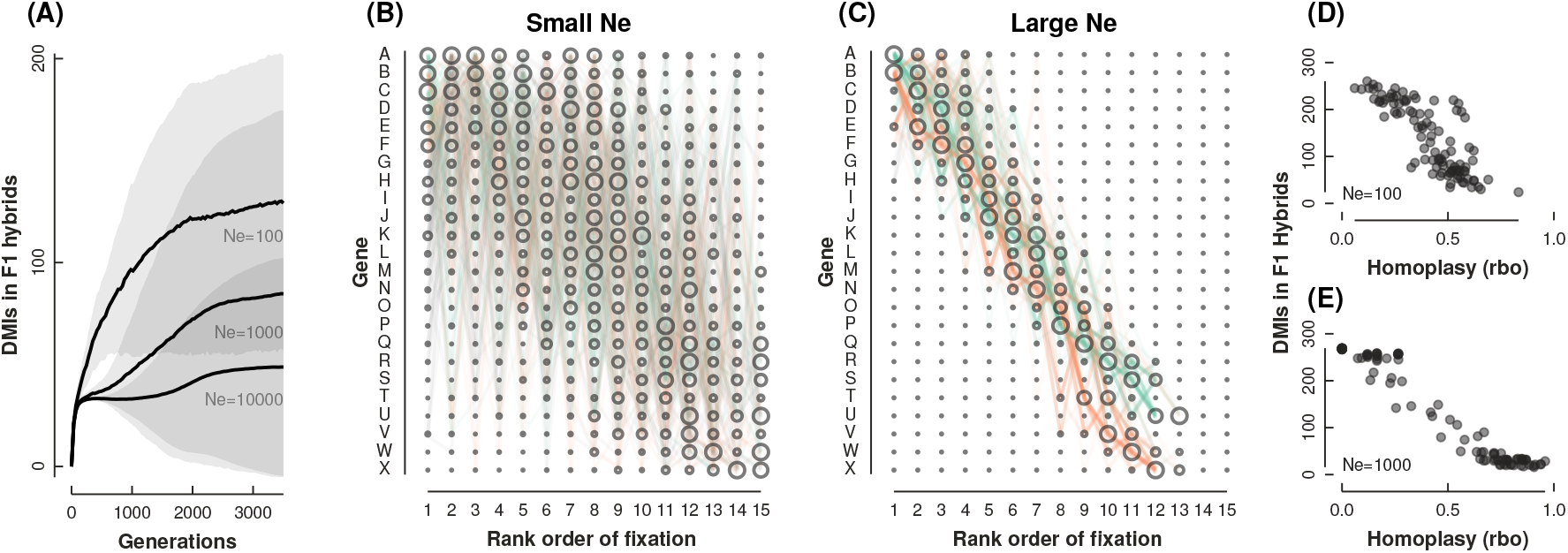
The impact of effective population size on the rate of DMI evolution between two populations under the individual-based (demographic) model with selection. Each figure shows variation across 100 replicate simulations. (A) Populations with small Ne evolve DMIs (mean±standard deviation) faster than populations with large Ne. (B-C) The order of fixation of new alleles across replicate simulations using the same set of pre-defined DMIs, where two optimal paths exist starting from either A (green) or B (orange). The size of circles represents the frequency with which an allele became fixed in a given rank order. (B) When Ne is small the fixation-order path is highly stochastic and many negative epistatic interactions become fixed within lineages. (C) When Ne is large purifying selection biases the order of fixation to follow optimal paths, minimizing fitness costs. (D-E) When populations evolve similar fixation-order paths they exhibit higher rates of homoplasy (measured as the rank-biased overlap of fixation orders) leading to lower rates of DMI accumulation in F1 hybrids.

The high variance in DMI accumulation rates under the Ind+Sel model reflects the high stochasticity in whether two lineages follow similar fixation order paths after diverging from a common ancestor. Simulations under a range of Ne values show that populations with small Ne are more likely to fix slightly deleterious mutations, leading to evolution along more highly stochastic fixation order paths (Fig. 3B). In contrast, populations with large Ne are more likely to evolve along one or more optimal paths that minimize the fixation of alleles with negative fitness consequences (incompatibilities) (Fig. 3C). When two populations evolve on highly stochastic fixation order paths they are more likely to fix many differences that contribute to many DMIs in their hybrids (Fig. 3B,D), whereas if both populations evolve under strong purifying selection, when Ne is larger, they are more likely to evolve similar fixation order paths (Fig. 3C) leading to greater homoplasy and fewer DMIs in F1 hybrids (Fig. 3E). Demography thus modulates the evolution of DMIs by allowing purifying selection to act on standing variation within populations, where epistatic interactions can expose fitness differences among different fixation order paths. When Ne is large this can bias populations towards evolving similar fixation orders, causing fewer DMIs in their hybrids.

### Empirical evidence for the demographic speciation model

For decades evolutionary theory has largely accepted that most mutations are deleterious and that genetic drift and purifying selection are more common than positive selection (*27*). Yet, over this same period positive selection has maintained prominence within speciation theory as the primary mechanism for the origin of species (*3, 24, 28*). The paucity of evidence for alternative mechanisms may reflect ascertainment bias in our typical search for few loci of large effect among already diverged species, as opposed to many loci of small effect among diverging populations with small ecological differences (*29*). Methods aimed specifically at detecting more subtle evidence of genetic incompatibilities, such as by analyzing genotype ratio distortions, support the view that genetic incompatibilities can occur widespread within species (*30*) – as predicted by our model.

A major strength of the demographic speciation model is that it provides a testable hypothesis for distinguishing the influence of positive selection (ecological speciation or traditional mutation-order speciation) from negative purifying selection (demographic speciation) in driving speciation. Our model predicts that lineages with smaller effective population sizes will accumulate DMIs more quickly, while populations with large Ne are more likely to maintain similar ancestral epistatic interactions and accumulate fewer DMIs. By contrast, ecological speciation makes the opposite prediction: populations with larger Ne should fix alleles under positive selection more quickly, and thus accumulate DMIs faster.

To test this prediction we fit a phylogenetic linear regression to the relationship between speciation rates (as a surrogate for the rate of DMI evolution) and geographic range size (as a surrogate for Ne) in two large-scale empirical datasets, representing approximately 10K species of birds (*5*) and 5K species of mammals (*31*). Because we expect the relationships between Ne and the rate of DMI evolution to exhibit high variance, we use surrogate measurements that are more widely available across massive-scale datasets, providing a unique opportunity to detect patterns only observable across many data points. Both datasets support a negative relationship (P<=0.006; Fig. 4) regardless of whether a phylogenetic correction is applied. Geographically widespread species (assumed to have large Ne) tend to have lower speciation rates (assumed to have slow rate of DMI evolution), whereas species with restricted geographic ranges (small Ne) exhibit higher speciation rates, as predicted by our simulations under the demographic speciation model.

**Figure 4:**
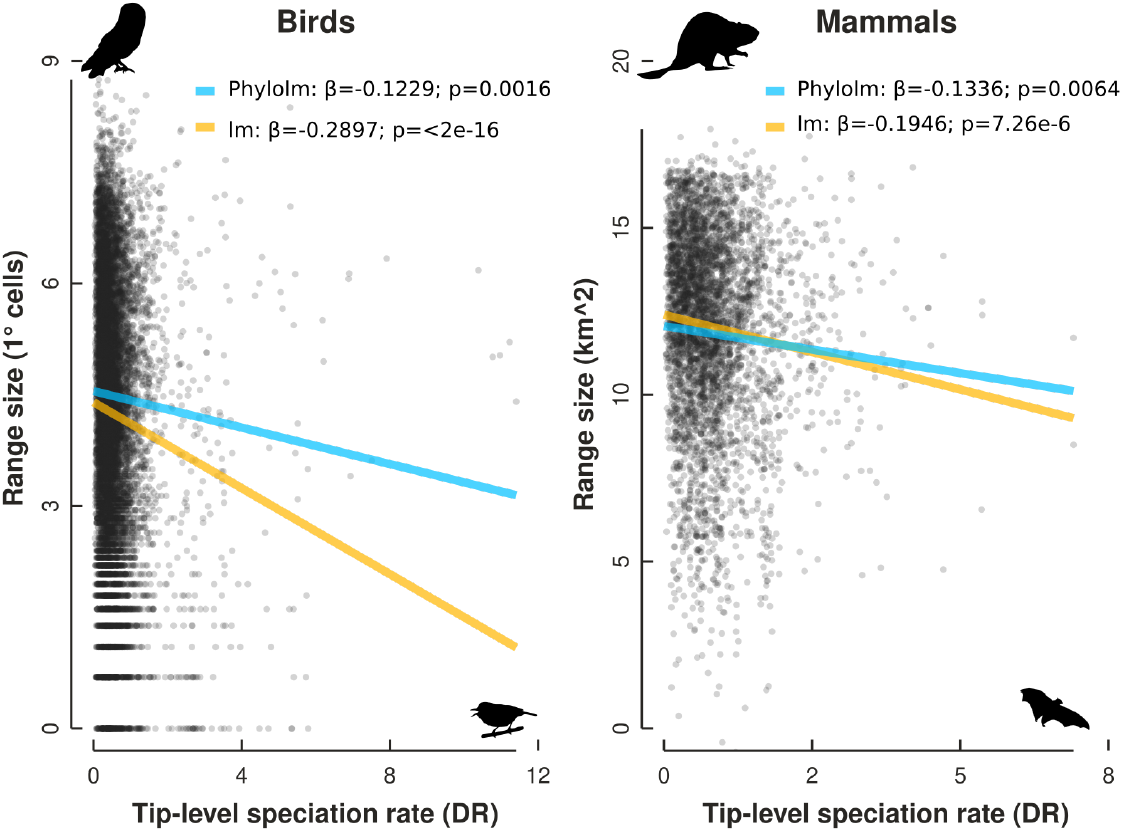
Linear regression between tip-level speciation rates (DR statistic) and geographic range size for species of birds (n=9,947) and mammals (n=5,422). The orange line is ordinary least squares, and the blue line phylogenetic least squares. Each point represents a species, and a silhouette (PhyloPic.org) is shown for a species from each extreme of the distribution.

### The Demographic Speciation Model

The demographic speciation model described here is a logical outcome of examining the evolution of epistatic interactions in the context of a population demographic model. We show that the faster-than-linear rate of accumulation of DMIs proposed in the snowball theory is a result of modeling a mutational process that does not allow for homoplasy. We also show that homoplasy is precisely the type of pattern purifying selection acting within a demographic population is likely to produce. Simulations under our new model demonstrate a simple process by which demography and purifying selection interact to affect the rate of accumulation of DMIs between lineages.

The demographic speciation model shares distinct similarities with the traditional mutation-order speciation process (*23, 24*), which defines the evolution of reproductive isolation as a chance outcome of the fixation of different alleles in two populations adapting by positive selection to similar selection pressures. Although our model also predicts that mutation order will be random, it differs from this model in predicting that fixation probabilities will be nonrandom, since selection can favor one outcome over another when the fitness effects of interacting alleles are exposed in the standing genetic variation within large populations (e.g., Fig. 1C). Interestingly, the best evidence for mutation-order speciation involves intragenomic conflicts, such as meiotic drive or cytoplasmic male sterility (*4,32*), where the fixation of a selfish element initially reduces overall fitness, and often occurs within genomic compartments with reduced effective population sizes, such as sex-linked chromosomes or plastids. The same is likely true for small-effect DMIs accumulating according to the demographic speciation process, where populations are more likely to accumulate genetic incompatibilities in genomic compartments with reduced effective populations sizes. A greater focus on identifying small-effect genetic incompatibilities within populations (*30*), and particularly by comparing large versus small isolated populations, may provide the best opportunities to investigate the impacts of demography on rates of evolution of DMIs.

Although the demographic speciation model emphasizes the effect of negative purifying selection, it does not preclude positive selection from occurring simultaneously. In fact, the very process of purifying selection acting on some allelic combinations in a population provides a mechanism for positive selection on other alleles, since individuals lacking incompatibilities may have a higher *relative* fitness. As an example, consider if “C” had been the first allele to become fixed in Fig. 1C, even though it causes the “Ca” incompatibility. This would impose positive selection on any new mutations that disrupt this incompatibility. If a derived “A” allele were to arise subsequently, for example, then individuals harboring this allele would have a higher relative fitness, since “AC” is not incompatible. The framework for such “distorter” and “restorer” type interactions (*24*) to evolve is inherent in our simulations simply by applying purifying selection. The potential for purifying selection to bias populations towards evolving similar fixation orders depends on the rate of fixation of new alleles (Fig. S3, Video. S1). In the case of very strong positive selection new alleles may fix too rapidly to interact with other standing variants that have not yet arisen, and the potential for purifying selection to bias populations towards evolving along fewer fixation order paths will be reduced.

Empirical studies have identified a disconnect between the rate at which reproductive isolating mechanisms evolve and the strength of postzygotic genetic incompatibilities (*6, 18*). The high variance in rates of evolution of genetic incompatibilities predicted under our model is concordant with these results, while also providing a framework for predicting why some lineages evolve DMIs faster than others. It may also provide new insights into the evolution of reproductive isolation by other mechanisms involving postyzgotic incompatibility. An example is reinforcement, which is only expected to occur when genes affecting assortative mating are in linkage disequilibrium with genes affecting postzygotic incompatibility (*33*). Under the perspective that DMIs involve few genes of large effect, reinforcement is expected to be rare, whereas under the view of our model, in which small-effect DMIs are abundant throughout the genome, reinforcing selection should occur frequently. In this way, the demographic speciation model, despite not modeling positive selection on phenotypes explicitly, can be reconciled with the diversity of reproductive variation that is abundantly evident in nature. Our new model challenges the prevailing views in speciation genetics rooted in positive selection, showing that instead purifying selection and demography together may play a major role in mediating the tempo and mode of speciation.

## Materials and Methods

### Simulating epistasis and DMIs

To demonstrate the effect of demography on the rate of accumulation of DMIs between diverging lineages we developed an individual-based forward-in-time simulation framework to model the evolution of negative epistatic interactions through time, using the software SLiM3 (*25*) (see General simulation framework in Supplementary Material and Methods). We modeled two populations that diverge from a common ancestor at time 0 and evolve independently for T generations, with F1 hybrids sampled periodically throughout the simulation that have no effect on the two parental populations (Fig. S4). Each population is composed of N diploid individuals that are initially homozygous for the ancestral allele at all sites. The genome consists of a single linear chromosome with 24 genes each 300 positions in length and 2000 positions apart. Genes are treated as a two-state character (ancestral or derived), and we explore a range of mutational models for which a mutation to any site within a gene causes a state transition (see Derived States and Reversibility in Supplementary Material and Methods).

We define a set of DMIs at the beginning of each simulation that is meant to represent constraints of the ancestral network of gene interactions that both populations inherit from their shared common ancestor. This set of DMIs is used to quantify incompatibilties in F1 hybrids, and to impose fitness costs to parental individuals harboring incompatibilities during simulations that include selection (Fig. S5) (see Quantification and Definition of DMI in Supplementary Material and Methods). The set of DMIs varies in each simulation, and is defined by a stochastic process following a series of rules that produces the maximum possible number of DMIs that could evolve for N genes under an infinite-sites mutational model (see Generating DMI sets/paths in Supplementary Material and Methods). A genome with 24 genes can evolve 276 DMIs. These rules ensure that there will always exist at least two fixation order paths (e.g., A fixes first, then D, then F) by which every gene could fix to the derived state in one population or the other without either population fixing an incompatible gene pair within it (Fig. S6). Each gene has the same number of possible incompatibilities, but varies in whether they involve DDI or ADI interactions with other genes. Consequently, the derived state of one gene may have zero incompatibilities with the ancestral genome, while the derived state of another gene has many incompatibilities with it, and is therefore unlikely to increase in frequency unless many other genes mutate first.

We explored seven simulation settings that vary in one or more assumptions that differentiate the Orr model from our theoretical ”demographic speciation model”, which is represented in our simulation study as the ”individual-based model with selection” (Ind+Sel) (see Assumptions and models in Supplementary Material and Methods). These assumptions include: (*1*) Instant fixation (Inst.), which causes new mutations to fix instantly across all individuals in a population; (*2*) Infinite sites (Inf.), which causes all mutations to occur at unique sites in both lineages; (*3*) Selection (Sel.), which imposes a quantitative fitness cost to parental individuals, scaled by the proportion of total DMIs contained in their genomes (in the absence of selection individuals may contain DMIs with no effect on their fitness); and (*4*) reversibility (Rev.), which causes sequential mutations to the same gene to revert its state back and forth between derived and ancestral states. Only a subset of all possible models could be tested as several parameter combinations give rise to conflicting assumptions. We performed a range of tests to validate that our results were independent of the effects of genome size (Fig. S7) (see Effects of genome size in Supplementary Material and Methods). To investigate the influence of effective population size variation on the accumulation of DMIs we ran simulations for the Ind+Sel model using different Ne values (100, 1000, and 10000), with 100 independent runs each. Finally, to quantify homoplasy in the order of fixation of derived alleles between parental lineages (Fig. 3), we ran three different simulations using the same predefined set of DMIs for each simulation. The “rank-biased overlap statistic”, a metric for comparing ranked lists, was used to measure the similarity in the order by which alleles fixed in each lineage in each simulation (see Quantifying homoplasy in fixation orders in Supplementary Materials and Methods). Detailed descriptions of the simulation process and reproducible code are available in the Simulation workflow and parameters section in Supplementary Materials and Methods.

### Testing demographic speciation model empirically

We used species-level phylogenetic trees and geographic range size datasets to measure the relationship between speciation rates and geographic range sizes. We used a posterior set of time-calibrated phylogenetic trees for 5,422 species of mammals (*31*), and for 9,947 bird species (*5*). Mammal geographic range sizes were extracted from the PHYLACINE database (*42*) and bird range sizes from the Open Traits Network database (*43*). Tip-level diversification rates (DR) were quantified following (*5*). Regression models were fit using linear models with or without phylogenetic corrections (see Demographic speciation model test in birds and mammals in Supplementary Materials and Methods).

## Supporting Online Material

Appendix I

Supplementary Materials and Methods

Figs. S1 to S8

Tables S1 to S2

References (*34–45*)

## Acknowledgments

This work was supported by an NSF grant to D.E. (DEB-1557059), and Columbia University startup funds. Thanks to B. Haller for advice on improving SLiM3 code, and to feedback on earlier versions of this manuscript from Tom Stewart, Trevor Price, and members of the Eaton lab.

## Appendix 1: The demographic speciation model

### Formal description

- The demographic speciation model (DSM) is a simplified evolutionary framework for making predictions about the evolution of intrinsic post-zygotic genetic incompatibilities between two or more independently evolving diverged lineages.

### Scope

- Predictions of the DSM can extend beyond the evolution of intrinsic post-zygotic incompatibilities to include additional evolutionary processes that may act in conjunction with, or as an outcome of, its core predictions. An example includes the evolution of premating reproductive isolation as a consequence of reinforcing selection against the formation of hybrids that exhibit low fitness due to the accumulation of DMIs under the DSM model.

### Definitions

- The Genome is modeled as a linear chromosome with genes separated by neutral regions in which recombination can occur.
- Genes are modeled as a binary character that can be in the ancestral or derived state. A mutation causes a transition to the derived state, and any subsequent mutations have no effect. More complex substitution models can be incorporated; a reversible binary state model was shown to have little impact on results.
- Genetic incompatibilities are modeled as two-locus epistatic interactions following the Dobhzansky-Muller model. When acting within parental lineages these are referred to as genetic incompatibilities, and in hybrids as genetic incompatibilities or DMIs.
- The fitness effect of a genetic incompatibility is the same whether occurring in a parent or a hybrid (e.g., 1%).
- The fitness effects of individual genetic incompatibilities are modeled as being drawn from a statistical distribution. In the paper we use a uniform distribution (e.g., 1%), but could similarly have used a normal normal distribution (e.g., mean=2%, stdev=0.5%).
- Although the fitness effects of individual genetic incompatibilities are drawn randomly from a statistical distribution, the fitness effects of individual alleles can vary greatly, since some will express many genetic incompatibilities, while others express few or none. The fitness effect of an allele is the additive effect of all of its genetic incompatibilities.
- The set of potential genetic incompatibilities (DMIs) is modeled as an inherited property of sister lineages when they diverge from a common ancestor. This set must contain at least two paths by which every gene could be mutated from the ancestral to a derived state in an order that does not give rise to any genetic incompatibilities. See the algorithm described in the paper for further details on how this is accomplished using infinite-sites mutations. Every gene has the same number of possible DMIs.
- Purifying selection is modeled as a probabilistic sampling process each generation based inversely on the proportion of total DMIs expressed in a diploid genotype (its fitness).

### Assumptions

- DMIs have small effects.
- Populations inherit the same gene interaction network (set of potential DMIs) at the time of divergence.
- The set of DMIs does not evolve, i.e., that diverged lineages continue to share the same set of ancestral DMIs after their divergence.

### Properties

- N lineages
- Ne population sizes
- G genes
- D two-locus DMIs defined by pairs of genes with one or both in the derived state.

### Expectations

- New alleles arise frequently but are typically lost due to genetic drift or purifying selection.
- Most new mutations are deleterious since they cause genetic incompatibilities with the gene interaction network. But alleles vary in the number of genetic incompatibilities they express, and thus in their fitness consequences. Alleles causing zero or few incompatibilities are neutral or nearly neutral and can become fixed by genetic drift in large populations.
- In large effective populations the order of fixation of new alleles is non-random. Populations are statistically biased towards evolving along fixation order paths that minimize genetic incompatibilities.
- Because two lineages inherit the same network of gene interactions they also inherit the same set of potential DMIs from their shared ancestor. This leads to statistically nonindependence between the lineages in their evolutionary trajectories. Each is biased towards evolving along ordered paths in the gene network that minimize genetic incompatibilities.
- When Ne is greater populations are more likely to evolve along similar paths that are closer to the optimum. This increases the probability of homoplasy, which decreases the number of DMIs that will evolve between species.
- Because lineages vary in effective population size the DSM can explain dramatic differences in rates of evolution of intrinsic post-zygotic genetic incompatibility even among closely related lineages.
- Populations with small effective populations sizes will accumulate DMIs more rapidly because they are likely to fix many alleles by genetic drift, despite their minor deleterious effects, leading to many alleles becoming fixed between species that cause many incompatibilities in hybrids (snowball-like).

## Supplementary Materials and Methods

### General simulation framework

Forward-in-time genetic simulations were performed using the software SLiM3 (*25*) following a Wright-Fisher process in which N diploid individuals are generated each generation by randomly drawing two haploid gametes from the previous generation. For this purpose we created a script called dmiSim.slim, written in the EIDOS programing language. This script takes different commands to toggle various features of the simulation and is available with full documentation of parameters at https://github.com/camayal/DMISimulations. We implement a simple two-population individual-based model to represent demography where two populations diverge from a common ancestor at generation 1 and maintain constant population sizes thereafter. At determined sampling periods a hybrid population is formed from which diploid F1 hybrid genotypes are sampled to calculate the number of DMIs that have accumulated. In the framework of SLiM3, this is accomplished by creating a new population of migrants drawn equally from both parental populations. During mating of this hybrid population individuals are suppressed if both of their parents are from the same population so that only F1 hybrids are created. The hybrid genotypes are saved to a file and the hybrid population is then discarded. The simulation then continues for the two parental populations until the next sampling period.

While a simpler simulation framework could have been developed for the questions addressed in this study, our aim was to establish an adaptable and extendable framework for studying the evolution of DMIs under arbitrarily complex models involving demographic parameters. By using SLiM3 we take advantage of a well tested codebase that also provides a framework to extend our models in the future to include more populations, geographic variation, quantitative traits, and other interesting components.

### Quantification and Definition of DMI

In the terminology of SLiM3 we define two regions in the simulated chromosomes: genes and spacers. Mutations in both occur at a per-site per-generation mutation rate of 1 × 10^−7^. In the context of our model, once a gene region mutates it becomes derived relative to the ancestral initial population, regardless of the number of subsequent mutations that may occur to sites within the gene region (except in models with reversibility). This information is used by our script to genotype each individual in each population to detect the presence of negative epistatic interactions in parental populations, or in F1 hybrids (where negative epistatic interactions are considered DMIs).

Negative epistatic interactions may occur between two alleles that are both derived (derived-derived; DDI) or between a derived and an ancestral allele (ancestral-derived; ADI) (*34*). As an example, if a diploid individual has the following genotype on two chromosome copies for five alleles (ABCdE, abCdE) and the following incompatibilities are defined (ADI: Da, Ea, Ed, Cb; DDI: AB, DB, CA, CD, CE, EB), then the diploid individual has 7/10 possible DMIs (Ea, Ed, Cb, AB, CA, CE, EB), three of which are DDI (Ea, Ed, Cb), four of which are ADI (AB, CA, CE, EB). Of these, six are trans effecting (between chromosome copies) and seven are cis effecting (same chromosome copy). A DMI can exist as both cis and trans and is counted as a single DMI.

### Derived States and Reversibility

The definition for a locus as being in a derived versus ancestral state is subjective relative to the starting point of the simulation. In this study the ancestral genome was always completely homozygous for an ancestral genotype. If a locus that is in the ancestral state (e.g., ”b”) becomes mutated within an individual in one of the parental populations, then it is considered to be in a derived state (e.g., ”B”). If this same gene mutates again in a different individual in the same or a different generation, it is also considered to have mutated to the same derived state (e.g., ”B”). If an individual that is already in the derived state experiences a mutation to the same gene again it will either remain in the derived state (”B”) or revert to the ancestral state (”b”) if the model includes reversibility.

This interpretation of genes as a binary state is a drastic over simplification; however, if we consider our primary interest to be the disruption of an ancestral gene network that is inherited by two populations, then the state of a gene can be interpreted in the context of either retaining or mutating its ancestral interactions with other genes. In this way, many derived changes may disrupt the ancestral interaction, and it does not matter whether they involve the exact same nucleotide substitutions, or whether the change creates a new function. For example, a DDI such as ‘AB’ can be interpreted as two genes with redundant functionality: “either ‘aB’ or ‘Ab’ would leave a functioning ancestral form, but ‘AB’ imposes a fitness cost because both gene functions have changed”. Or more simply, “if the ancestral function ‘a’ is lost then ‘b’ must be present, and vice versa”. Similarly, an ADI such as ‘aC’ can be interpreted like a suppressor gene: “if the ancestral function of ‘c’ is lost then ‘a’ cannot be present”.

For this study, we simulated divergence of two populations from a common ancestor that is completely fixed for ancestral alleles, and therefore we do not model the effects of standing variation inherited from a common ancestor. The inclusion of standing ancestral variation is likely to be important for accurately modeling the rate of accumulation of DMIs in empirical studies (*35*), since the amount of sequence divergence between two populations at time 0 will be greater than 0. In the context of our simulations it is simpler to disallow ancestral variation and thus we can fit our regression analyses as starting with zero DMIs at time zero. In the future our simulation framework could be expanded to incorporate ancestral polymorphisms which would allow for interesting examinations of models with more than two populations to examine how the inheritance of standing variation in a phylogenetic context affects patterns of DMI accumulation (*36*).

### Assumptions and models

We evaluated the assumptions of the Orr (1995) mathematical model by testing different simulation models that relax its main assumptions independently (e.g., instant fixation and infinite sites). In addition, we introduce several new models by toggling two additional assumptions: reversibility and selection. We introduce reversibility and selection as potentially important factors for their ability to maintain sites in an ancestral state, which can significantly affect the accumulation of ancestral-derived type DMIs. In total seven models were tested that include different combinations of compatible assumptions (Table S1). Some combinations were not tested either because they are impossible in combination, such as infinite sites and reversibility, or because they are computationally difficult together, like demographic scenarios and infinite sites.

We applied selection following a “multiple incompatibility” approach (*37*), where individual DMIs each express the same quantitative effect on fitness, and multiple DMIs cumulatively can cause complete inviability. Selection applies to the two parental populations during the formation of offspring. Gametes are randomly sampled to form new diploid offspring which either survive and contribute to the population in the following generation, or die and are discarded. If an offspring is discarded a new diploid offspring is sampled until the new population reaches the set population size. The probability that an offspring survives to join the new population depends on its fitness which is determined by the number of negative epistatic interactions in its genome. This affect is applied proportional to the total number of DMIs defined in the simulation. For example, in a simulation with 100 possible DMIs, if an individual harbors 2 DMIs it has 98% probability of survival; if it harbors 98 DMIs then it has a 2% probability of survival.

Selection only applies to individuals in the parental populations. F1 hybrids do not experience selection since they are removed from the simulation after DMIs are counted and do not affect the evolution of the parental lineages.

Our individual-based model differs from Orr (*10*) in both the inclusion of demography and in its mutational considerations. In terms of demography, the population size in Orr’s model is effectively one, since new mutations fix instantaneously. By contrast, new mutations in the individual-based models arise in a single haplotype and subsequently evolve by drift (or additionally by selection) such that variation can persist within populations over many generations.

In terms of mutational processes, Orr’s model assumes an infinite sites model in which mutations never occur in the same gene in both lineages, or revert to an ancestral state. It should be noted that this special case of infinite-sites applies to both populations simultaneously, such that they are effectively not evolving independently, since a mutation in one lineage prevents it from occurring in the other. By contrast, our individual-based model allows that both lineages could mutate the same gene multiple times, and a mutation in one lineage has no effect on the evolution of the other lineage (Fig. S4).

To demonstrate that our results with respect to Orr’s original model are not a mere consequence of using simulation versus mathematical modeling, we included a modification of our general SLiM3 simulation framework to simulate an approximation of Orr’s model that includes the infinite sites and instantaneous fixation assumptions. Modifications required to emulate Orr’s mathematical model include: (*1*) a random mutation was drawn each generation and applied to all individuals in one population or the other (i.e., instant fixation); (*2*) hybrid populations are sampled every generation when a substitution occurs; (*3*) mutation rate was zero to avoid random mutations in the genome that are different than the instant-fixation we apply in step 1; and (*4*) the number of generations is set to three times the number of genes simulated (because two additional generations are needed to complete migration and hybrid production as explained above). In order to evaluate the effect of the infinite-sites assumption on the Orr model we also implemented an option to turn-off this assumption (Orr-inf) by allowing for multiple mutations to occur probabilistically to any gene each generation, with the possibility that the same genes can mutate to a derived state in both lineages. This probabilistic test is repeated each generation until all genes mutate to a derived state in both lineages. Each model is named according to its base assumption (Orr vs Individual-based). We then append additional modifiers representing deviations from each model (e.g., Orr-Inf, Orr-Inf+Rev, Ind+Sel, Ind+Sel+Rev).

### Generating DMI sets/paths

We defined a set of DMIs that could evolve for a set of N genes prior to the start of each simulation using a stochastic process that follows a set of defined rules, described below. A Python program, dmiGenerator.py (https://github.com/camayal/dmiGenerator), was written to automate this task. It takes as input either a user-supplied genotype for N genes (e.g., “AbCD”), or if not provided, it generates a random genotype for N genes where each gene is sampled as a random binomial to be either ancestral or derived. From this genotype it then returns a set of all two-locus gene interactions that could occur in a perfect snowball scenario where one lineage evolves this genotype and the other lineage evolves the exact opposite (i.e., infinite-sites; Fig. S6).

For example, we can provide as input the gene sequence “AbCD” and the program will return the set of all incompatibilities for 4 genes that could exist while allowing a population to evolve from “abcd” to “AbCD” without ever fixing an incompatibility: (“AB”, “CB”, “DB”, “Ca”, “Da”, “Dc”). The incompatibilities are generated by evolving a mutational path between the two genomes following a set of rules based on Orr’s theory that prevents any DMIs from evolving within lineages while maximizing the number of DMIs that evolve between two lineages. This includes: (*1*) a mutation cannot evolve that is incompatible with other genes in the same genome (*10*); (*2*) a derived allele cannot be incompatible with its own alternative allelic state (e.g., derived and ancestral) (*9*); (*3*) incompatibilities are only considered between populations; and (*4*) incompatibilities are asymmetric (*2*).

It is important to note that the process used here to define the set of DMIs for individual simulations is distinct from whether those DMIs actually evolve during a simulation. The set of DMIs itself can be thought of as the constraints imposed by the ancestral network of gene interactions that both populations inherit from their common ancestor. There exists one or more ordered paths of fixation through this network whereby every gene can be mutated and fixed without experiencing any fitness cost. However, most mutations to this network will disrupt some part of it and thus impose a fitness consequence. Mutations are random and can occur in any order. But whether or not a population evolves along the optimal ”fixation order path” will depend on the strength of purifying selection to prevent the fixation of deleterious mutations.

The output of dmiGenerator.py is piped into our SLiM3 script, dmiSim.slim, to perform simulations. The results of each simulation are reported in two JSON files, one that stores the F1 hybrid genotypes at each sampling interval (infoHybs.json), and another which contain the number of DMIs in each sampled F1 hybrid cohort (numHybsByNumofDMIs.json).

### Simulation workflow and parameters

For each simulation model 100 replicate runs were performed from different random starting seeds. Simulations were run for a variable number of generations depending on the model type. Orr models require few generations (70) to finish since the infinite sites and instant fixation assumptions lead to a steady-state result rapidly. The Orr model with reversibility took longer to reach a steady state (1500 generations) since multiple mutations can yield more variable results. Neutral individual-based models (Ind) were run for at least 250,000 generations to ensure simulations reached a steady state, while models with selection (e.g., Ind+Sel) were run twice as long, which appears to generally capture a plateauing rate. In the much longer running individual-based models F1 hybrid cohorts were generated and saved to a file at uneven intervals (6,250 - 25,000 generations) to sample more densely at the beginning of simulations when patterns change more rapidly. DMIs in hybrid populations were counted in the SLiM script and plots were generated using Python scripts (extractAverageDMis.py and multiple-TimeGraphs.py).

We explored a range of simulation parameters to identify settings that would allow simulations to finish in a reasonable amount of time even under complex models. Running time increases for larger population sizes and numbers of genes. We explored values for populations sizes in (10, 100, 500, 1K) and numbers of genes in (*6, 12, 24*). In the absence of demography population size has no effect on either run times or results, so we used *N_e_* = 2. Neutral individual-based models take longer to run when population sizes are larger, but results are not qualitatively affected by population size (Fig. S2), so we used *N_e_* = 100.

For models that include selection and multiple individuals (demography) both run times and results are affected by the population size, and we report results for different population sizes (Fig. 3). The per-site per-generation mutation rate (*μ* =1 × 10^−7^) and recombination rate (*r* = 1 × 10^−8^) were kept constant across all simulations except for those used in Fig. 3. In this case, rates were down-scaled to (μ = 1 × 10^−5^) and (r =1 × 10^−6^) in order to increase the speed of simulations with very large effective population sizes. Population sizes and the number of generations were scaled proportionally.

All code, including slurm batch scripts for distributing simulations in parallel on a cluster, are available on GitHub (https://github.com/camayal/DMISimulations).

### AIC model fit to DMI accumulation curves

To evaluate the best model fit to rates of DMI accumulation through time under each simulation model we compared the following functions: linear *ax* + *b*, quadratic *ax^2^* + *bx* + *c*, or logistic 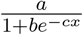. A script was written in R to perform model fitting (AICw.R) using nonlinear least squares analysis in the ’nlsLM’ function implemented in the ’minpack.lm’ package (*38*). AIC model weights were computed with AICcmodavg (*39*) to compare the model fit of each function for each model. Models were also fit to the accumulation curves for separate classes of DMIs (ADI and DDI; Table S2) following the same methodology using R code in (AICw_DDDA.R).

### Effects of genome size

We investigated whether the finite genome size used in our simulations (e.g., 24 genes) influenced the slowing patterns of DMI accumulation observed across all models except the Orr model. For this, we examined the effect of removing simulations from our results that reach either the maximum number of DMIs possible (e.g., 276) or several lower values (>200, >250, >270). Despite removing these ”saturating” scenarios the trend of a logistic pattern of DMI accumulation is still observed (Fig. S7), suggesting that genome size saturation is not driving our results.

### Zombie effect

Simulating DMIs under an individual-based model introduces new complications that are not observed in the Orr model, where the infinite sites and instant fixation assumptions ensure that every new mutation has the potential to create one or more DMIs. In that case, DMIs can simply be calculated as each new mutation arises without the need to define ahead of time which gene interactions from a finite sized genome will or will not cause incompatibilities. In contrast, we have found that this problem cannot be ignored in individual-based models.

This is clear from a simple example: if two derived mutations occur in one population (AB), this could create an ancestral-derived incompatibility (ADI) in which aB bears a fitness cost; however, the other population may have already experienced this allele combination in an earlier generation before it was defined as a DMI (Fig. S5). This problem is particularly relevant when selection against DMIs is included in the model, since aB should have been selected against in an earlier generation in population 2, but the DMI was not yet defined until a particular sequence of mutations occurred later in population 1. We refer to this sequence-dependent effect on DMIs as the ”zombie effect”. By defining a set of all possible DMIs from an ancestral genome at the beginning of a simulation, and selecting a subset that is not contradictory, this problem is easily avoided, although the combinatorics of calculating DMIs can become quickly burdensome for very large numbers of genes. Our simulations involve relatively small genomes (24 genes) but are large enough that we can clearly distinguish the rate of accumulation of DMIs from patterns that are affected by saturation of the genome where most or all genes are converted to the derived state.

### Quantifying homoplasy in fixation orders

We quantified homoplasy in the order of fixation of derived alleles between two lineages using the ’rank-biased overlap’ statistic, a measure of similarity between ragged, possibly infinite ranked lists which may or may not contain the same items (*40*). Code was implemented from the rbo.py script (https://github.com/dlukes/rbo).

This measure importantly reflects not only whether two lineages evolved similar genotypes at the end of a simulation, but also whether the differences among them arose early versus later in their sequence of fixations. This is of interest since when two populations fix the same mutations early after divergence they are more likely to fix similar subsequent mutations, due to the epistatic effects of the earlier fixed alleles on the fitness of subsequent mutations. This is demonstrated in our results by the strong correlation between rbo and the number of DMIs (Fig. 3). When two lineages share 50% of genes in the same derived or ancestral state, the identify of those genes determines whether they cause few or many DMIs.

A complete snowball scenario, as observed in the Orr model, only occurs due to the infinite-sites assumption where each lineage evolves the exact opposite set of derived genes. Our results show that this is unlikely to occur in an individual-based model since selection will favor similar fixation order paths that minimize the evolution of negative epistatic interaction within lineages. In other words, the greater similarity of the fixation order paths between two populations the less likely they are to experience a snowballing pattern of incompatibilities.

### Demographic speciation model test in birds and mammals

We examined large-scale bird and mammal datasets to test the unique prediction of the demographic speciation model of a negative correlation between effective population size (Ne) and the rate of DMI evolution. Because both Ne and the number of DMIs are poorly quantified for most taxa, we used surrogate measurements that are expected to be highly correlated. Geographic range size was used as a proxy for effective population size (*41*), and speciation rates estimated from time-scaled phylogenies was used as a proxy for the rate of DMI evolution.

Geographic range sizes were extracted from global databases on mammal or bird life history traits. For mammals we used range size estimates from the PHYLACINE database (*42*) recorded in units of square kilometers. For birds we used estimates from the Open Traits Network database (*43*) recorded in units of 1 degree grid cells.

Speciation rates were calculated from posterior distributions of time-calibrated trees using the tip-level diversification rate statistic (DR) as described in (*5*), and implemented in the toytree Python package (*44*). Phylogenetic trees for mammals and birds come from the vertlife datasets (*5, 31*). Linear regression models were fit in R using the ‘lm‘ package, or ‘phylolm‘ to include phylogenetic correction (*45*). The formula used for both regression tests is: log(Geographic range) ~ Diversification rate.

---

**Figure S1.**
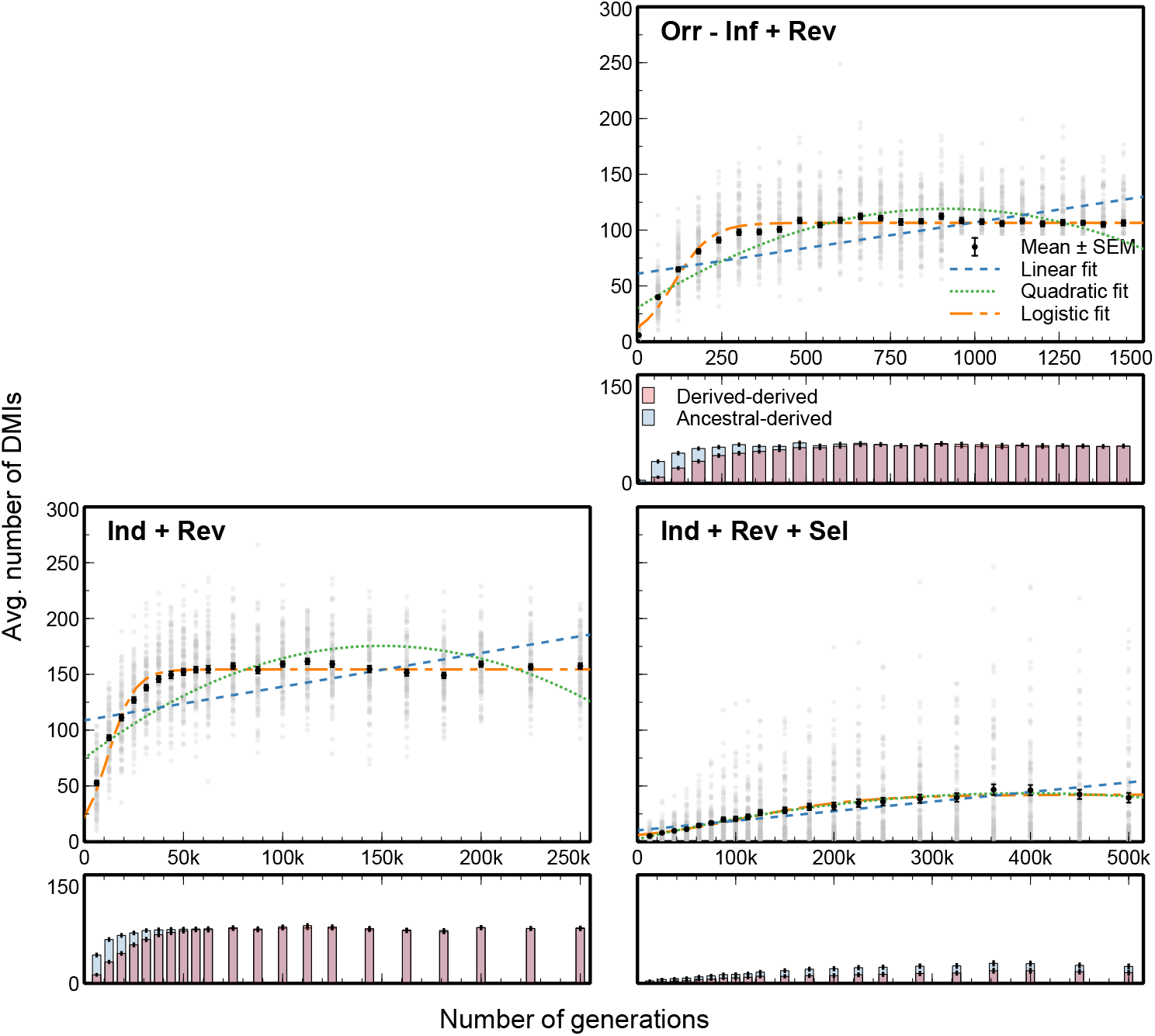
The rate of accumulation of epistatic gene interactions causing Dobzhansky-Muller incompatibilities (DMIs) under several additional simulation that include reversibility in the mutation process. Upper panels show the average number of DMIs (black) in hybrid cohorts with standard error bars and gray points representing individual simulation results. Colored curves show the best fit regression model with parameters fit by nonlinear least squares analysis using three different functions (blue dashed for linear, green dotted for quadratic, and orange dash-dotted for logistic). Lower panels show DDIs and ADIs with standard errors for each model. Reversibility acts to maintain many alleles in their ancestral state which increases the frequency of ADI.

**Figure S2.**
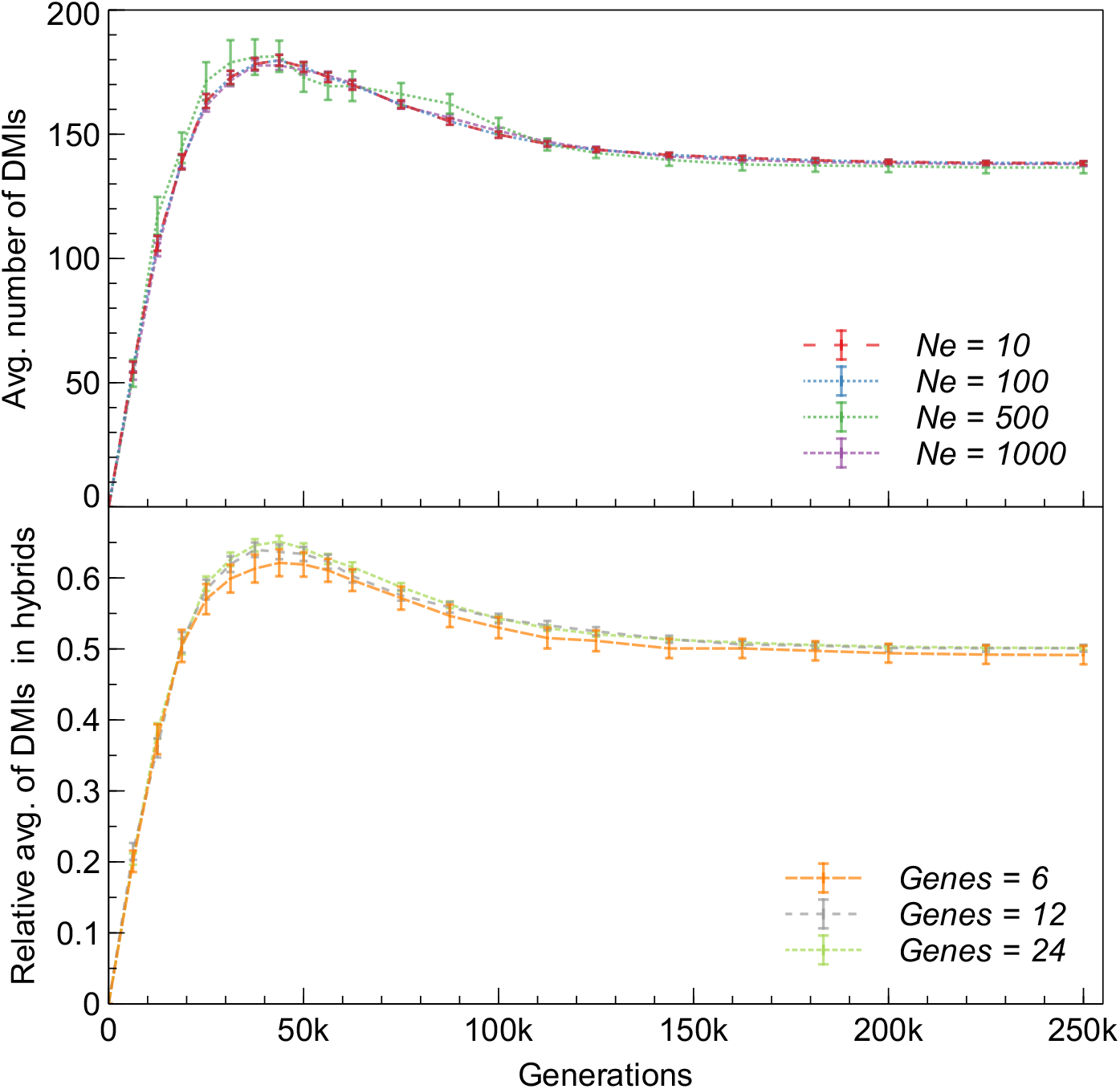
Effective population size (*N_e_*) and the number of genes (genome size) have no effect on the rate of accumulation of DMIs under neutral individual-based model simulations.

**Figure S3.**
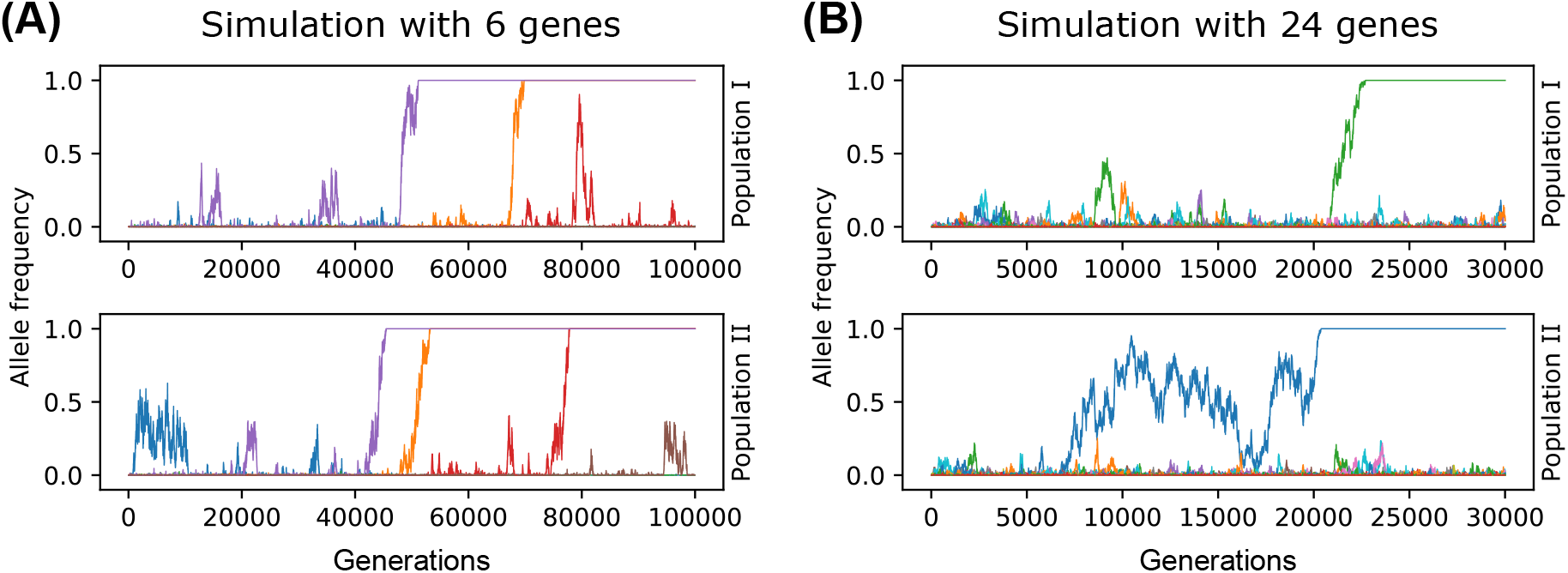
Demography and selection caused by epistatic interactions among standing genetic variation within populations affects the fixation probabilities of alleles in the Ind+Sel model. (A) A simulation with six genes where both lineages initially fix the same two alleles in the same order. The red allele becomes unconstrained by purifying selection only after the first two alleles become fixed. It is now neutral, and becomes fixed in one population before the end of the simulation, but not in the other, simply due to genetic drift. (B) Simulations with 24 genes involve a more complex network of epistatic interactions, where many new mutations arise but exhibit low fitness early on and are suppressed by purifying selection.

**Figure S4.**
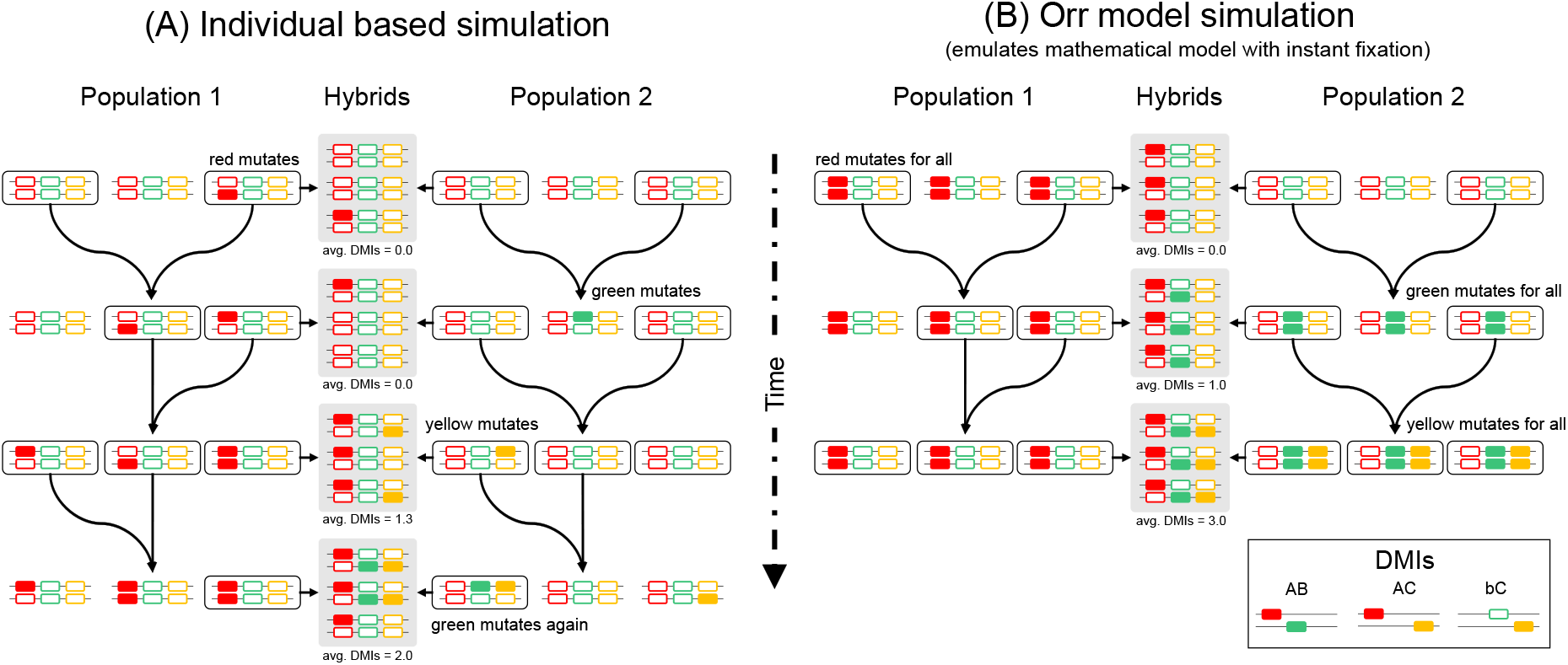
Our individual-based simulation (A) differs in several key ways from the mathematical framework used by Orr (B) to demonstrate the snowball theory. Most notably, new alleles arise in single haplotypes in the individual-based simulation, versus being fixed across all individuals instantly in the Orr model. This slows the rate of accumulation of DMIs because new alleles may be lost by drift (e.g., green allele) or take time to reach fixation (e.g., red allele). When alleles occur at low frequencies they may not contribute to the number of DMIs counted in a sampled hybrid population simply due to sampling effects.

**Figure S5.**
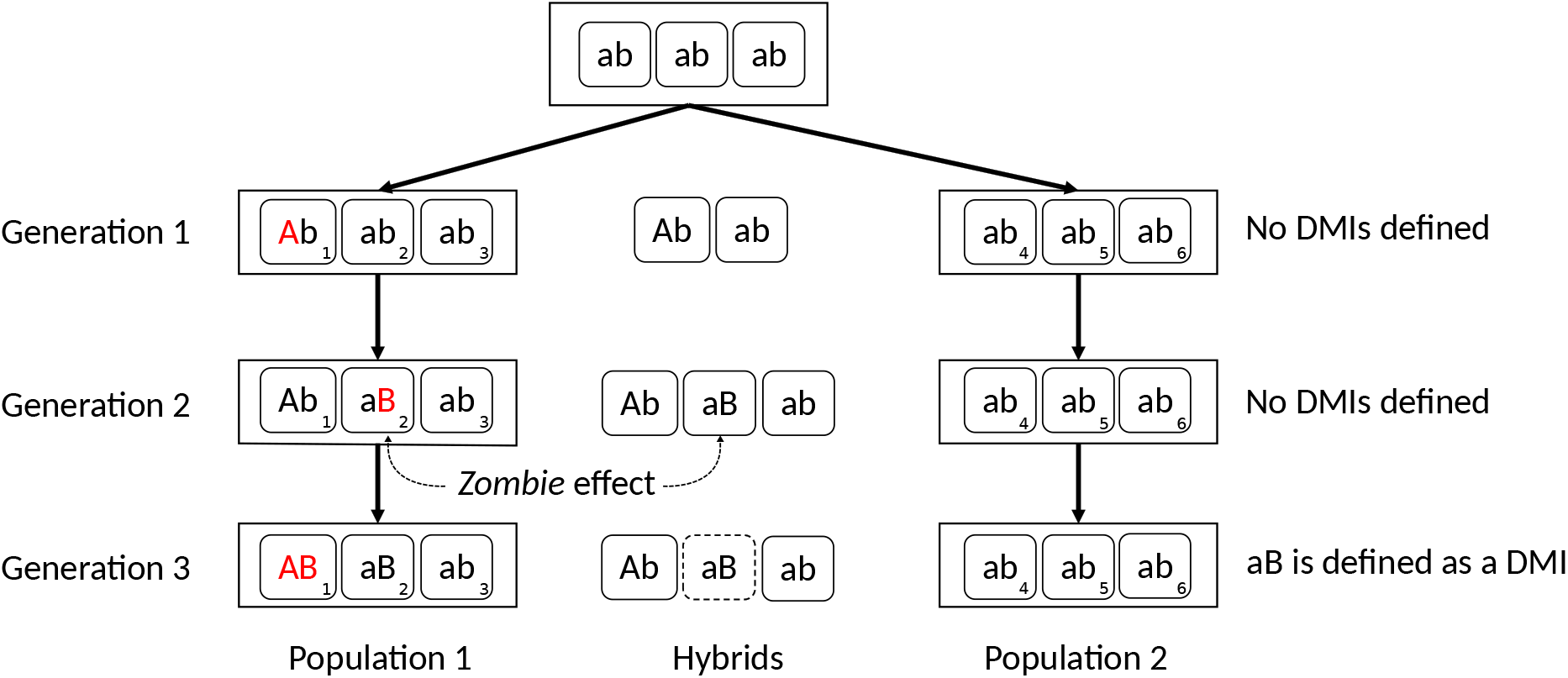
Schematic representation of an example of the *zombie effect* that can arise if epistatic interactions causing DMIs are not pre-defined before starting a simulation. A mutation ”A” arises in an individual in generation 1, and a different mutation ”B” arises in another individual in generation 2. At this point following the definition of a DMI neither ”A” nor ”B” should be incompatible with an ancestral allele. In generation 3 recombination brings the two derived alleles together in the same genome to form ”AB”. Upon observation of this genotype we could define an ancestral-derived incompatibility such as ”aB” under the assumption that ’a’ has not been tested with the ”B” allele in the same genome. In the traditional Orr model the ”aB” DMI would be defined in generation 3. However, in a individual-based model like this example the genotype ”aB” has already occurred in generation 2, both in population 1 and in the hybrid cohort. To properly apply purifying selection against negative epistatic interactions that can evolve within populations it is thus necessary to define all DMIs before the beginning of the simulation.

**Figure S6.**
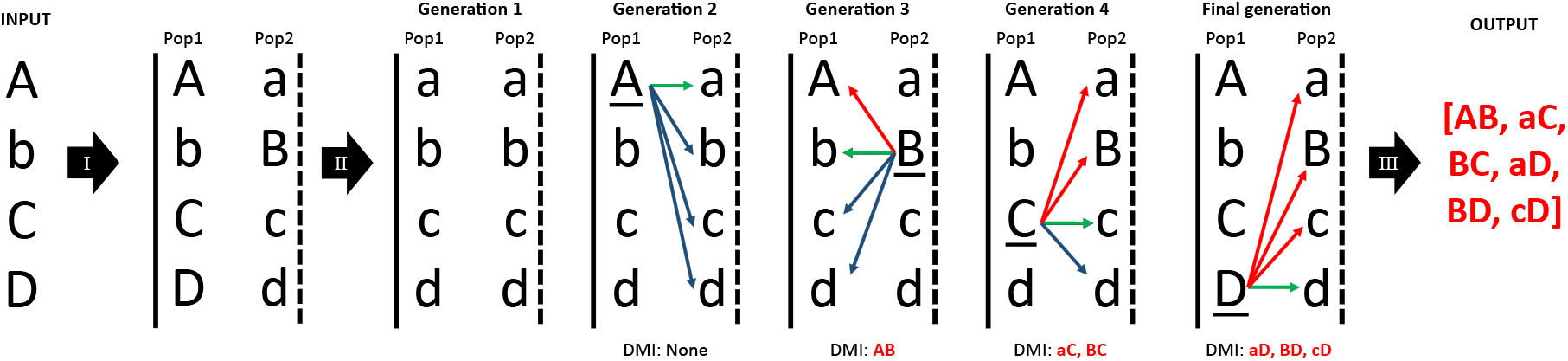
Every simulation begins by sampling a set of two-locus DMIs that represents epistatic constraints imposed by the ancestral network of gene interactions inherited by both populations. The tool dmiGenerator.py generates the set of DMIs following a defined logic. It takes as input an ordered sequence of allelic states for N genes, where upper or lower case represents derived versus ancestral states, respectively. The DMI generating logic applies in three steps: (I) the input sequence is assigned as the end-state for one population and the opposite sequence is assigned as the end-state of the other. (II) Both populations then start from the ancestral genome and fix one mutation at a time, in the order of the input sequence. As each gene mutates the genomes of the two populations are compared to identify whether any new valid DMIs have formed. DMIs cannot occur between two allelic states of the same gene (green arrows), or between allelic states that occur together in the genome that experienced the new mutation (blue arrows). Other gene combinations (red) represent possible genetic incompatibilities under the snowball model. This leads to a set of DMIs where the derived state for some genes is incompatible with many ancestral alleles, while the derived state of other genes may be incompatible with few or no ancestral alleles. (III) The cycle of mutations and comparisons between genomes is continued until all genes in the input sequence have been visited sequentially, and the list of DMIs is returned.

**Figure S7.**
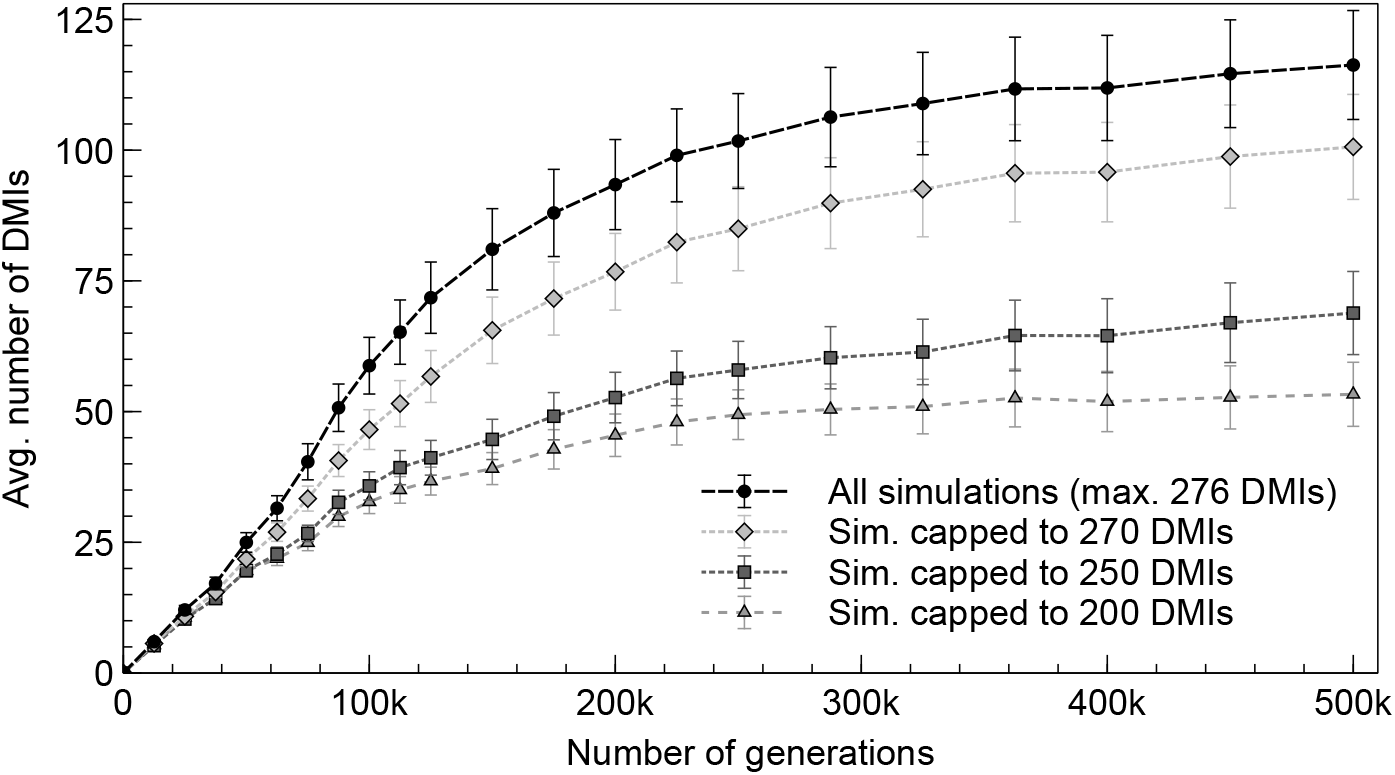
Finite genome size limits do not cause the slowing-down pattern of DMI accumulation observed in all models except the Orr model. This is demonstrated by removing all simulation replicates that fixed all possible DMIs (276) or a set lower limit of DMIs.

**Video S1.** A video showing replicate simulations from different random seeds for a demographic + selection model. For ease of visualization this model includes only four genes. The top and bottom panels show the allele frequencies of the two diverged populations through time, and the middle panel the allele frequencies of hybrid populations. In most simulations the two populations fix alleles in a similar order due to purifying selection. When the two populations fix different alleles in a different (e.g., the final simulation shown) hybrid genotypes contain derived and ancestral forms of all genes (allele frequencies near 0.5) leading to greater incompatibilities.

**Table S1.** All possible models that can be simulated with the four model assumptions explored in this study. Some are types are impossible to simulate and others present computational difficulties or unrealistic scenarios.

| Assumptions |  |  |  | ID | Model name | Conflicts |  |  | Conflict | Tested |
| --- | --- | --- | --- | --- | --- | --- | --- | --- | --- | --- |
| Inst. | Inf. | Sel. | Rev. |  |  | Ind+Inf | Orr+Sel | Inf+Rev |  |  |
| 1 | 1 | 0 | 0 | 1100 | Orr |  |  |  |  | Yes |
| 1 | 1 | 1 | 0 | 1110 | Orr+Sel |  | 1 |  | Yes |  |
| 1 | 0 | 0 | 0 | 1000 | Orr-Inf |  |  |  |  | Yes |
| 1 | 1 | 0 | 1 | 1101 | Orr+Rev |  |  | 1 | Yes |  |
| 1 | 0 | 1 | 0 | 1010 | Orr-Inf+Sel |  | 1 |  | Yes |  |
| 1 | 0 | 0 | 1 | 1001 | Orr-Inf+Rev |  |  |  |  | Yes |
| 1 | 0 | 1 | 1 | 1011 | Orr-Inf+Rev+Sel |  | 1 |  | Yes |  |
| 1 | 1 | 1 | 1 | 1111 | Orr+Rev+Sel |  | 1 | 1 | Yes |  |
| 0 | 0 | 0 | 0 | 0000 | Ind |  |  |  |  | Yes |
| 0 | 0 | 1 | 0 | 0010 | Ind+Sel |  |  |  |  | Yes |
| 0 | 1 | 0 | 0 | 0100 | Ind+Inf | 1 |  |  | Yes |  |
| 0 | 1 | 0 | 1 | 0101 | Ind+Inf+Rev | 1 |  | 1 | Yes |  |
| 0 | 1 | 1 | 0 | 0110 | Ind+Inf+Sel | 1 |  |  | Yes |  |
| 0 | 1 | 1 | 1 | 0111 | Ind+Inf+Rev+Sel | 1 |  | 1 | Yes |  |
| 0 | 0 | 0 | 1 | 0001 | Ind+Rev |  |  |  |  | Yes |
| 0 | 0 | 1 | 1 | 0011 | Ind+Rev+Sel |  |  |  |  | Yes |

**Table S2.** Model comparisons using AIC to fit regression models to rates of accumulation of different types of DMIs through time.

| Model | AICwt |  |  |  |  |  |
| --- | --- | --- | --- | --- | --- | --- |
|  | Derived-derived |  |  | Ancestral-derived |  |  |
|  | Linear | Quadratic | Logistic | Linear | Quadratic | Logistic |
| Orr | 0 | 1 | 0 | 0.59 | 0.41 | 0 |
| Orr-Inf | 0 | 0 | 1 | 0.02 | 0.01 | 0.07 |
| Orr-Inf+Rev | 0 | 0 | 1 | 0 | 0 | 1 |
| Dem | 0 | 0 | 1 | 0 | 0 | 1 |
| Dem+Rev | 0 | 0 | 1 | 0 | 0 | 1 |
| Dem+Sel | 0 | 0.92 | 0.08 | 0 | 0 | 1 |
| Dem+Rev+Sel | 0 | 1 | 0 | 0 | 1 | 0 |

